# High-channel-count neural recording and stimulation platform with 5,376 simultaneous recording channels

**DOI:** 10.64898/2026.03.13.709972

**Authors:** Yingying Fan, Yuhang Ma, Pavlo Zolotavin, Gerald Topalli, Weinan Wang, Mattias Karlsson, Magnus Karlsson, Lan Luan, Chong Xie, Taiyun Chi

## Abstract

Advancing neural interfaces requires large-scale, high-density recording technologies capable of capturing full-spectrum neural activity across cortical and subcortical regions. Here, we present a scalable approach to integrate neural electrodes with advanced application-specific integrated circuits (ASICs). Specifically, we custom-designed an ASIC with 5,376 simultaneous channels, each sampling at 20 kS/s and enabling >1.3 Gb/s total data streaming throughput. The ASIC incorporates in-pixel amplification, time-division multiplexed ADCs, and on-chip stimulation capabilities, ensuring precise signal acquisition with minimal power consumption while maintaining a low noise level of 5.5 µVrms. We further developed an interconnect strategy using gold bump bonding, which allows for high-density integration of the flexible probe and rigid chip. We demonstrate the capacity of this platform through the integration with a flexible μECoG array. The resulting device allows for the high-resolution mapping of *in vivo* field potentials on the cortical surfaces of rat brains, supported by the precise localization of evoked sensory activities. These results prove an effective approach towards highly integrated neural interfaces with applications in brain-computer interfaces, neuroprosthetics, and large-scale functional brain mapping.

## Introduction

There is a growing interest in the neuroengineering community in developing technologies capable of recording from large populations of neurons across both cortical and subcortical regions^1–14^, driven by both scientific and translational applications. Scientifically, large-scale, high-density recordings enable functional mapping of distributed brain areas, which reveals the underlying circuits of sensorimotor processing, decision-making, and memory formation ^4,15–21^. On the translational side, increasing channel counts enhances the decoding accuracy of motor intentions and cognitive states for brain-machine interfaces ^17,22^, thereby improving the precision and reliability of prosthetic control and neural modulation systems ^23–25^.

A key technical bottleneck in scaling neural interface lies in the availability of high-channel-count backend electronics capable of amplifying, digitizing, and streaming signals from thousands of channels while simultaneously supporting programmable stimulation. Conventional implementations typically rely on off-the-shelf, moderate-density backends integrated with probes through custom packaging. For instance, systems have scaled to 1,024 channels using 16 commercially available 64-channel Utah arrays (Blackrock Microsystems) ^26,27^, and over 1,000 channels by pairing custom flexible probes with Intan electronics^14,28–30^. While accessible and modular, such an approach often leads to suboptimal system specifications with respect to power consumption, weight distribution, spatial arrangement, and heat dissipation. These issues become increasingly challenging for chronic and freely moving experiments ^31,32^.

Emerging approaches leverage application-specific integrated circuits (ASICs) to directly support high-density neural recording. Neuropixels 1.0 and 2.0, for example, offer 384 active channels selected from a total of 960 or 5,120 electrodes ^13,23^. However, due to hardware constraints, these systems require switching among different banks of electrodes, leaving most of the electrodes unused at any given time. Their rigid silicon probes also raise concerns about long-term biocompatibility^34^. Similarly, the Broadband Integrated Silicon Circuit (BISC) employs a ~50 µm-thick ASIC designed for μECoG integrating 65,536 electrodes^5^. It supports 256 channels at 33.9 kS/s or 1,024 channels at 8.475 kS/s for increased coverage, though the latter configuration is below the typical action potential (AP) sampling requirement.

In this work, we present a scalable neural recording and stimulation platform that overcomes these trade-offs through co-optimization of ASIC design, flexible electrode array fabrication, and packaging. The system features three key elements: (1) a compact ASIC with 5,376 simultaneous recording channels and a field-programmable gate array (FPGA) backend, capable of raw, full-bandwidth data acquisition (>1.3 Gb/s); (2) high-density, flexible µECoG electrode arrays for brain mapping in rodents and humans; and (3) a scalable, high-yield bonding method for integrating high-density electrode arrays with ASICs. We further validated this system through in vivo high-density µECoG recordings in rats.

### System overview

The complete system (Fig. □1a-d) is housed in a compact 3 □cm × 3.3 □cm × □1.2 □cm package. As illustrated in Fig. 1b, the headstage features a vertically stacked PCB design consisting of three layers: a top ASIC board, a middle routing board, and a bottom FPGA board. The ASIC board hosts the recording/stimulation ASIC and local decoupling capacitors. The routing board serves as an interposer, connecting the ASIC to the FPGA. The FPGA board interfaces with the main control unit (SpikeGadgets LLC, San Francisco, CA, USA) via a microHDMI cable, transmitting the digitized neural data for downstream storage and processing.

**Figure□1.**
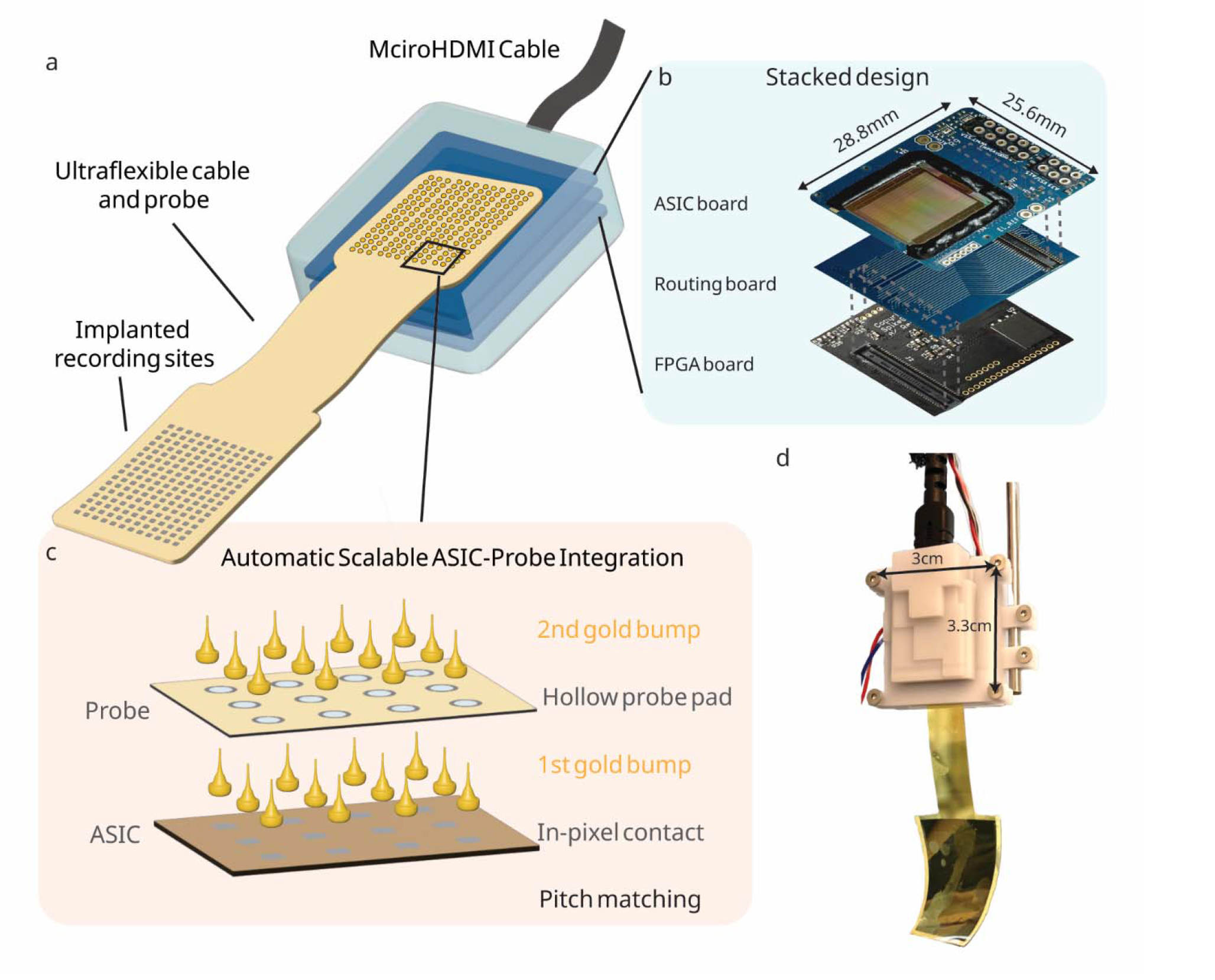
Overview of the 5,376-channel, full-band neural data acquisition system. **a** Illustration of the complete system. A lightweight headstage encloses three stacked PCBs. An ultra-flexible, cable-shaped neural probe is bonded at one end to the ASIC, while the other end contains the implanted recording sites interfacing with the cortex. A microHDMI cable connects the headstage to an external controller, enabling high-speed data streaming and ASIC control. **b** Exploded view of the stacked PCB design. From top to bottom: an ASIC board for recording and stimulation, an intermediate routing board, and an FPGA board for data interface control. **c** Illustration of the dual-bump ASIC–probe bonding process. First-layer gold bumps are deposited onto in-pixel contacts of the ASIC. The ultra-flexible probe, fabricated with hollow pad openings, is aligned and overlaid. Second-layer gold bumps are then formed to bridge the probe’s hollow-pad gold boundary and first-layer bumps, creating direct electrical and mechanical connections. This dual-bump scheme enables precise, one-to-one, pitch-matched bonding across all 5,376 channels, while preserving mechanical flexibility and supporting seamless connections between the ASIC and probe. **d** Photograph of a fully assembled headstage, showing the compact system with integrated flexible probe and microHDMI cable.

The ASIC employs a 2D scalable architecture (Fig. □1c), consisting of an 84 × 64 pixel array (5,376 pixels in total) with a pixel size of 180 □µm × 180 □µm. This pitch size was optimized to maximize density while supporting reliable bonding with a variety of neural probes, including both surface and penetrating probes^35^, making the system versatile across a wide range of neurophysiological applications. Each pixel contains a dedicated low-noise analog front-end that amplifies neural signals from 2 Hz to 10 kHz.

The probe-ASIC bonding is achieved by a dual-bump bonding scheme (Fig. □1c). Before bonding, first-layer Au bumps (Ø □≈ □70 □µm) are plated directly onto each in-pixel pad of the ASIC. Then, the ultraflexible probe, patterned with hollow-pad openings (outer Ø □≈ □85 □µm, inner Ø □≈ □60 □µm), is aligned over these bumps. Second-layer gold bumps are deposited into the hollow pads, fusing with the underlying first-layer bumps to form 1:1, pitch-matched electrical and mechanical connections. The resulting stack height is < 45µm. This bonding method avoids post-processing of CMOS chips, achieves >90% bonding yield, and maintains the mechanical compliance of the flexible probe. The entire bonding process for all 5,376 channels is completed within ~10 minutes (Supplementary Video 1 and 2) using a standard thermocompression ball bonder, offering a scalable, low-cost integration pathway for mass production.

### ASIC architecture and electrical benchmarks

The custom-designed ASIC integrates four key functions (Fig. □2a): 5,376-channel simultaneous recording, 224-channel programmable current stimulation, impedance measurement, and electrode electroplating (Methods). The central array consists of 84 □× □64 = 5,376 recording pixels, with each pixel incorporating a low-noise amplifier (LNA) and a programmable gain amplifier (PGA) (Supplementary Fig. S1). This per-pixel analog front-end architecture ensures optimal noise performance and efficiently utilizes the pixel area. Every four pixels share a 12-bit fourth-order ΣΔ analog-to-digital converter (ADC) through a 4:1 time-division multiplexer. The ADCs are placed outside the pixel array (Fig. 2b), similar to CMOS image sensor design ^36^. Each ADC operates at 80 □kSa/s, resulting in an effective sampling rate of 20 kSa/s per pixel, sufficient for digitizing AP signals. This specification was chosen to establish the ASIC as a universal backend capable of supporting various neural probe modalities.

**Figure□2.**
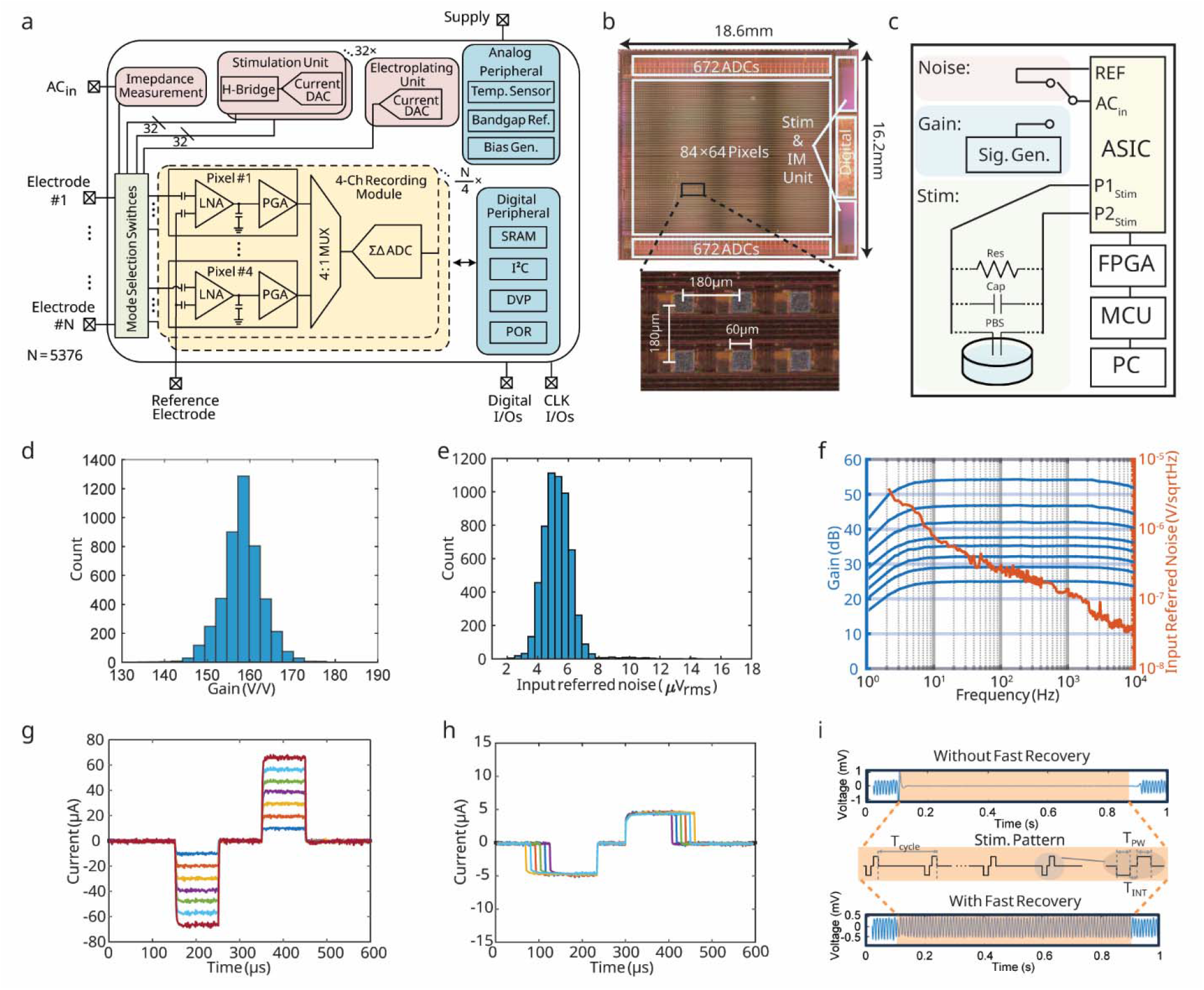
Architecture and performance of the 5□376-channel recording– stimulation ASIC. **a** Block diagram of the ASIC integrating four key functionalities: 5,376-channel simultaneous recording, 224-channel current stimulation, impedance measurement, and electrode electroplating, programmed by in-pixel mode selection switches. Every four pixels share a 12-bit ΣΔ ADC via a 4:1 time-division multiplexer, with an effective sampling rate of 20 □kSa/s per pixel. Digitized data are serialized and streamed off-chip through dual-lane digital video port (DVP) interfaces at 1.3 □Gb/s, while an I^2^C interface programs the ASIC. **b** Micrograph of the fabricated ASIC, highlighting the 84 □× □64 pixel array and surrounding peripheral circuitry. Zoomed-in inset shows individual pixel dimensions (180 □µm pitch). **c** Experimental setup for noise, gain, and stimulation measurements. **d** Histogram of midband gain across all 5,376 channels, showing excellent uniformity with a mean of 158.2 □V/V and a standard deviation of 3.2 V/V. **e** Histogram of integrated input-referred noise (2 □Hz–10 □kHz), with a mean of 5.3 □µV_rms_ and a standard deviation of 0.99 µV_rms_. **f** Measured frequency response from a representative pixel, showing eight programmable midband gains (25.2 □dB – 54 □dB, left axis) and the corresponding input-referred noise power spectral density (right axis), with 5.5 □µVrms integrated noise over the 2 □Hz–10 □kHz band. **g** Programmable stimulation currents ranging from 200 □nA to 70 □µA with 200 □nA resolution, demonstrated using a resistive load. **h** Biphasic current stimulation pulses with adjustable pulse widths. **i** Continuous neural recording during delivery of 50 □Hz biphasic stimulation with 10 □µA pulses, 167 □µs cathodic, 67 □µs inter-phase gap, 167 □µs anodic (T_period_ = 20 ms, T_PW_ = 167 µs, and T_INT_ = 67 µs). A fast recovery gate disables the front-end during the 0.7 □ms stimulation window, enabling the amplifier to regain full sensitivity and capture neural activity before the next stimulation cycle.

Peripheral circuit blocks surrounding the pixel array manage data streaming, clocking, and control. Digitized data are serialized by dual-lane Digital Video Port (DVP) interfaces (Supplementary Fig. S3), supporting an aggregate data rate of 1.3 Gb/s. An I^2^C interface programs the key parameters of the ASIC, such as voltage gain and filter cut-off frequency. It has an average power consumption of 42 µW per channel, leading to a power density of 0.75 mW/mm^2^ that remains safely within our thermal budget. As detailed in the Supplementary Information Sec. 1 and Supplementary Fig. S4, data transmission accounts for the majority of this power. Because the ASIC is integrated into the external headstage rather than being implanted intracranially, this thermal load is isolated from the cortical surface. The heat is effectively managed through natural convection and the thermal mass of the PCB, ensuring the implanted probe remains at body temperature. The power consumption can be reduced by setting inactive circuit blocks into standby. When full-channel recording is not required, selective channel activation proportionally scales power consumption, providing a flexible balance between power consumption and data throughput.

The ASIC demonstrates excellent channel-to-channel uniformity. Histograms of midband gain and integrated input-referred noise across the full array are shown in Fig. 2d–e. The midband gain distribution shows a mean of 158.2 V/V and a standard deviation of only 3.2 V/V. Input-referred noise, integrated over the 2 Hz–10 kHz band, shows a mean of 5.3 µVrms and a standard deviation of 0.99 µVrms. The analog front-end supports eight programmable midband gain settings (25.2 – 54.1 dB), with a corresponding integrated noise floor of 5.5 µVrms (Fig. 2f). This measurement confirms that the overall noise performance is dominated by the analog front-end, with minimal contribution from the ADC and digital interface.

The ASIC also supports high-fidelity programmable current stimulation. The stimulator supports programmable output from 200 nA to 70 µA in 200 nA steps, leveraging well-matched unit current sources for linearity and accuracy. Biphasic pulses were demonstrated with high accuracy under various loads (Fig. 2g-h, Supplementary Figure. S2). To support simultaneous stimulation and recording, a fast recovery mechanism was implemented. The front-end is briefly disabled during each stimulation pulse and regains full sensitivity within 1.5ms, enabling artifact-free neural signal acquisition before the next pulse (Fig. 2i).

### High-Density ECoG Probe Design and Electrochemical Characterization

To enable large-scale brain surface mapping, we developed flexible ECoG arrays with 5,376 channels, specifically designed for integration with the custom ASIC. Figure □3a shows the layout and fabricated results of two probe variants: one designed for human cortical recordings (left) and one optimized for rodent models (right). Both designs share a common backend architecture that incorporates 5,376 gold-coated hollow ring pads to facilitate reliable integration via dual gold bump bonding with the ASIC.

**Figure□3.**
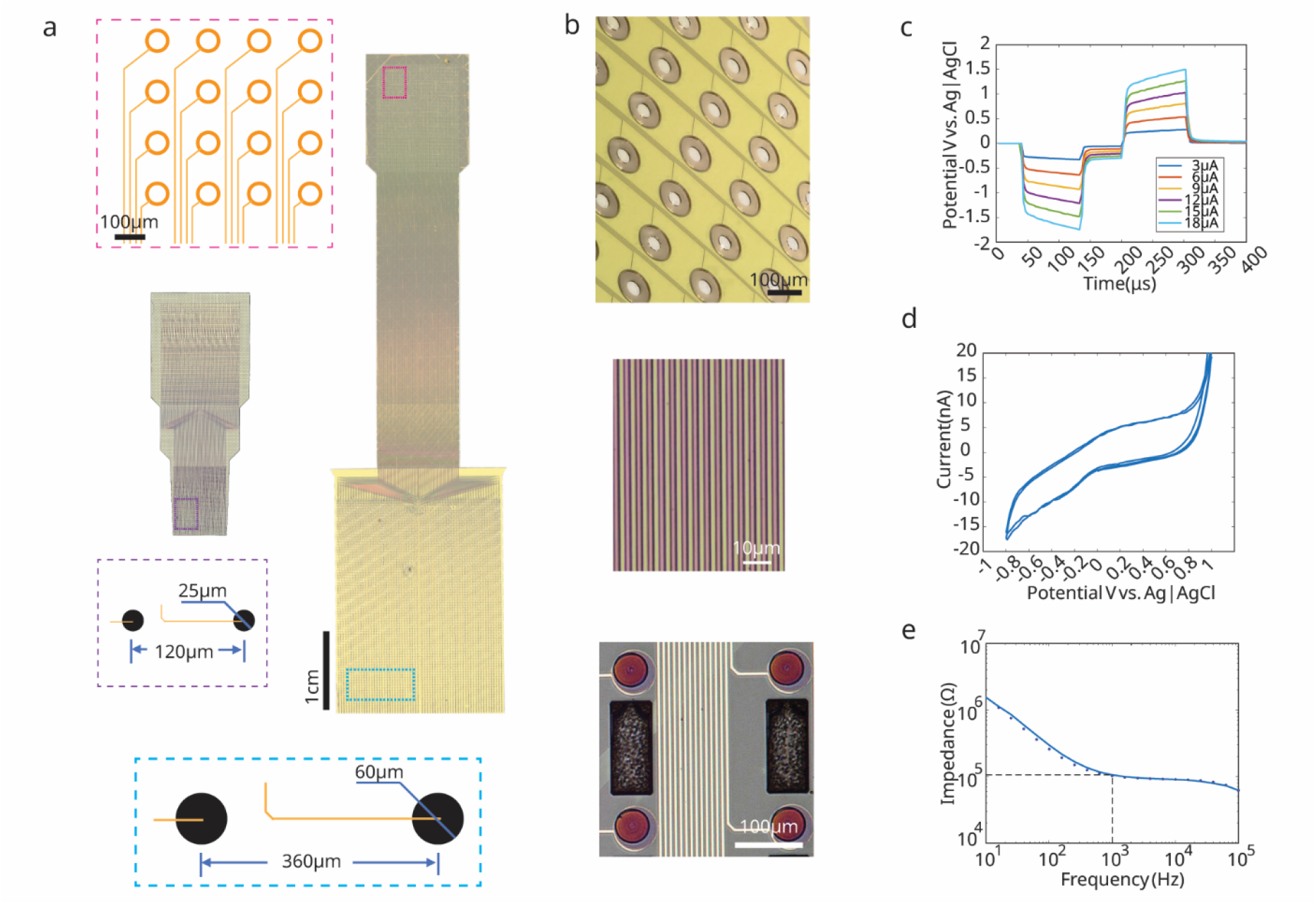
Probe design & characterization. **a** Photographs of two ultraflexible polyimide-based probes designed for electrocorticography (ECoG) applications in human (left) and rodent (right) models. Insets show zoomed-in layouts of the backend bonding pads (top left, dashed pink box) and recording electrodes (bottom, dashed purple and blue boxes). Electrode diameters range from 25 □µm to 60 □µm, with inter-electrode spacing tailored to different application scales. **b** (Top) Optical image of the hollow bonding pad structure designed for gold bump integration with the ASIC. (Middle) Close-up of the 4-layer gold interconnects separated by polyimide insulation layers, demonstrating dense routing capability. (Bottom) Image of an IrOx-coated electrode sites with a rectangular perfusion hole between them. Scale bar, 100 □µm. **c** Voltage transient measurements of the sputtered Iridium Oxide (IrOx) electrodes under varying stimulation currents (3–18 □µA) in phosphate-buffered saline (PBS). **d** Cyclic voltammetry (CV) curves of the IrOx electrodes, showing reversible redox behavior and confirming good charge injection capability. **e** Electrochemical impedance spectroscopy (EIS) of the IrOx electrodes, showing impedance magnitude as a function of frequency; typical impedance at 1 □kHz is ~100 □kΩ, suitable for low-noise neural recording.

Each probe consists of four metal routing layers, with 1,344 channels per layer. The multilayer design enables dense interconnects with a minimum metal linewidth of 2 □µm and spacing of 2.5 □µm. The middle portion of the probe is constructed using a polyimide cable, providing both mechanical flexibility and dense signal routing. In the rodent version, it minimizes the distance between the cortical surface and recording electronics, thereby improving signal integrity in freely behaving experiments.

The electrode layouts were tailored for species-specific neurophysiological requirements. For the human array, electrodes measure 60 □µm in diameter and are spaced 360 □µm apart, covering a 30.5 □mm × 22.9 □mm cortical area. This spacing ensures broad cortical coverage while avoiding signal contamination from spiking activity, making it suitable for capturing population-level dynamics across functional areas such as the temporal lobe (speech decoding), motor cortex (movement intention), and somatosensory cortex (tactile feedback). For the rodent design, the array spans 10 □mm × 7.6 □mm, with 25 □µm electrodes spaced 120 □µm apart, enabling high-resolution mapping of cortical field potentials over a wide hemispheric region.

Figure □3b highlights critical structural elements of the probe. The top panel shows the hollow bonding pads used for pitch-matched integration with the ASIC. The middle panel reveals the four-layer gold interconnect stack, separated by polyimide insulation layers to preserve flexibility and routing density. The bottom panel presents a close-up of a recording electrode coated with sputtered iridium oxide (IrOx), including a central perfusion opening to allow fluid exchange and improved tissue contact.

To evaluate electrochemical performance, we characterized the IrOx electrodes under conditions representative of both stimulation and recording. Voltage transient responses measured in phosphate-buffered saline (PBS) at varying current amplitudes (3–18 □µA) are shown in Fig. □3c, confirming consistent electrode behavior across the stimulation range. Cyclic voltammetry (CV) curves recorded at 100 □mV/s (Fig. □3d) demonstrate reversible redox behavior with high charge storage capacity, indicating suitability for safe and efficient stimulation. Electrochemical impedance spectroscopy (EIS), shown in Fig. □3e, reveals a characteristic low-impedance profile across frequency, with typical 1 □kHz impedance values of 101 □kΩ—ideal for high signal-to-noise ratio (SNR) neural recording.

### Scalable Dual-Bump Bonding and Integration Performance

To enable wafer-scale integration between the neural probe and ASIC, we adapted a MicroFlex-style interconnect ^37^ and extended it to a dual gold bump bonding strategy optimized for high channel density and mechanical robustness. Figure □4a–f illustrates the bonding process and resulting structures. First, gold bumps (~70 □µm diameter) were deposited onto the aluminum contact pads of the ASIC, forming the first-layer bump array across all 5,376 pixels (Fig. □4a). A zoomed-in view (Fig. □4b) confirms the uniform height and pitch of the bumps, and Fig. □4c shows the hemispherical profile of a representative bump. Each bump was fabricated with a tapered tail, which facilitates precise alignment of the ASIC and the probe in the next step.

**Figure 4.**
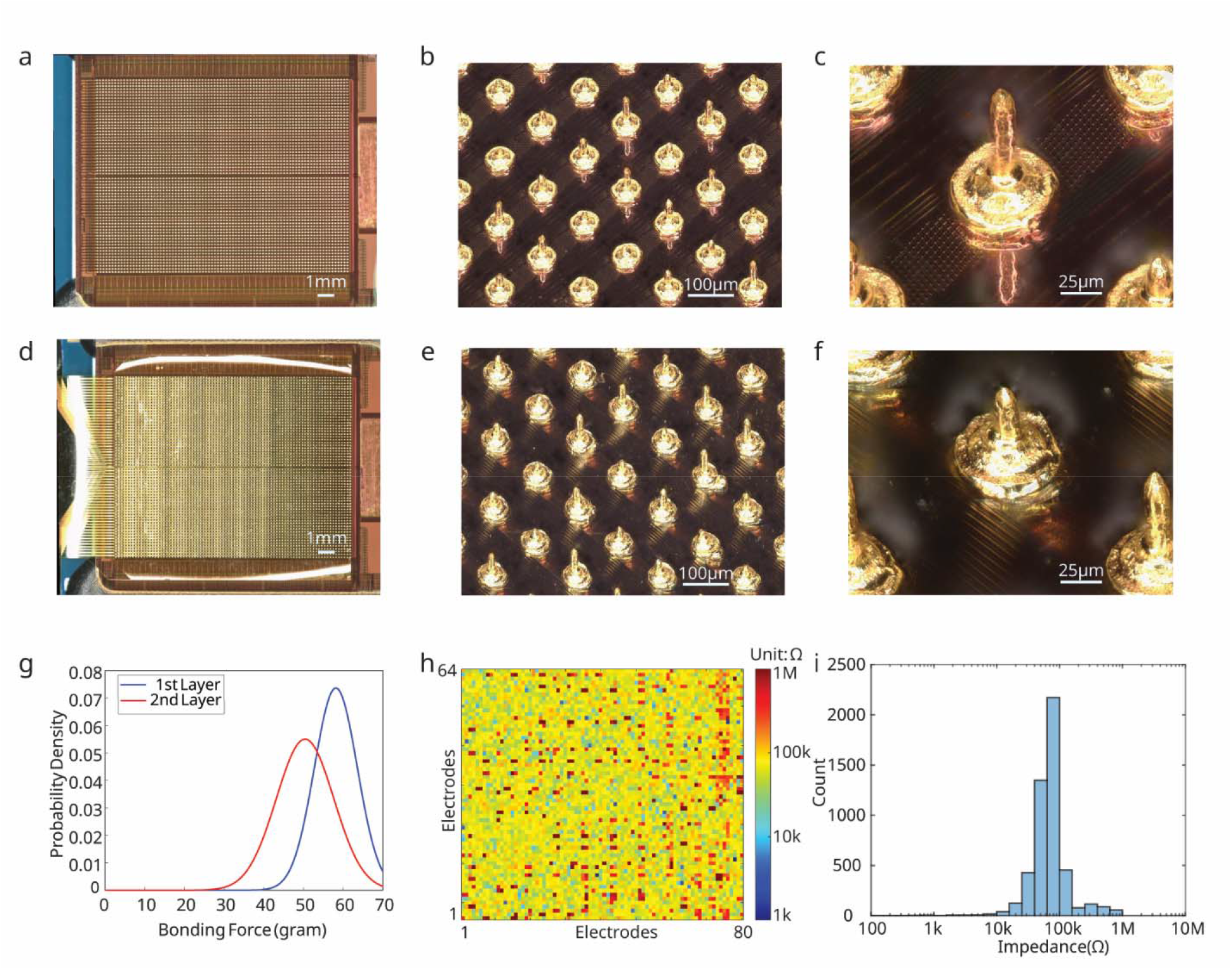
Dual gold bump bonding process and post-bonding electrical characterization. **a** Optical image of the ASIC with the first-layer gold bumps deposited onto each in-pixel aluminum pad across the full 5,376-pixel array. **b** Zoomed-in view of multiple first-layer bumps, showing uniform height and alignment. **c** Close-up image of a single first-layer bump, highlighting its dome-shaped profile. **d** Optical image of the bonded assembly after placement of the flexible ECoG probe and deposition of second-layer gold bumps through the probe’s hollow pads. **e** Zoomed-in view of multiple bonded bumps demonstrating uniform filling of the hollow pads. **f** Close-up image of a single fully bonded gold bump, showing complete fusion between the first and second layers. **g** Distribution of bonding force measured for first-layer (red) and second-layer (blue) gold bumps, indicating good mechanical consistency and successful formation of reliable interconnects. **h** Impedance mapping of all 5,376 electrodes in the ECoG array after bonding, measured in phosphate-buffered saline (PBS). Most connections show impedance values within the expected range. **i** Histogram of impedance values from panel h, centered around ~80 □kΩ at 1 □kHz, confirming high electrical yield across the array.

Next, the flexible ECoG probe, patterned with hollow ring-shaped bonding pads, was aligned and placed over the ASIC pixel array. A second-layer gold bump was deposited through each hollow via (Fig. □4d). These second-layer bumps serve a dual function: they fuse with the underlying first-layer gold bumps and simultaneously establish contact with the gold-coated ring pads on the probe, creating a vertical electrical and mechanical interconnect between each ASIC pixel and the corresponding probe electrode. Fig. □4e shows a zoomed-in view of multiple dual-bump junctions, and Fig. □4f provides a close-up of a well-formed gold-to-gold bond. The smooth, complete filling of the hollow pad by the second-layer bumps ensures low contact resistance and high reliability.

To assess bonding strength, we conducted destructive shear tests by applying a unidirectional force at 100 □µm/s until detachment occurred. As shown in Fig. □4g, the first-layer bonds (Al/Au interface) exhibited an average failure force of ~60 □g, while the second-layer bonds (Au/Au interface) failed at ~50 □g. The stronger adhesion at the Al/Au interface is attributed to the plasma treatment of the aluminum surface. The slightly lower shear force at the Au/Au interface is likely due to porosity in the evaporated gold and minor variation in contact uniformity.

Following successful bonding, the ASIC-probe module was fully assembled with the stacked supporting electronics, including the FPGA, routing board, and power supply. The bonding region was encapsulated using underfill epoxy and medical-grade silicone to provide mechanical reinforcement and environmental protection, enabling long-term stable operation during in vivo use.

We further evaluated the electrical yield of the bonded system through impedance spectroscopy in phosphate-buffered saline (PBS). As shown in Fig. □4h, a full impedance map across the array demonstrates uniform electrode performance. The corresponding histogram in Fig. □4i reveals that 92% of electrodes exhibited impedance below 100 □kΩ, with a prominent peak centered around 85 □kΩ. This range is ideal for capturing both local field potentials (LFPs) and multi-unit activity (MUA), balancing low noise with sufficient coupling to extracellular signals.

### In Vivo Cortical Mapping Using a 5,376-Channel μECoG Array

We demonstrated the functionality of our high-density μECoG platform through acute cortical mapping in adult rats. To minimize environmental noise including 60 □Hz interference, electromagnetic coupling, and ground loops, we implemented several noise-reduction strategies: (1) enclosing the recording setup in a grounded Faraday cage, (2) grounding the ASIC, neural probes, and reference independently, and (3) physically separating the reference and ground electrodes.

The flexible μECoG probe was placed onto the cortical surface, covering multiple functional areas including somatosensory, motor, and visual cortex (Fig. □5a,b). During recording, mechanical stimuli were delivered in separate trials to the whiskers, forelimb, hindlimb, and trunk. Stimulation events were time-stamped and aligned with the recorded neural signals streamed through the head-mounted system.

**Figure 5.**
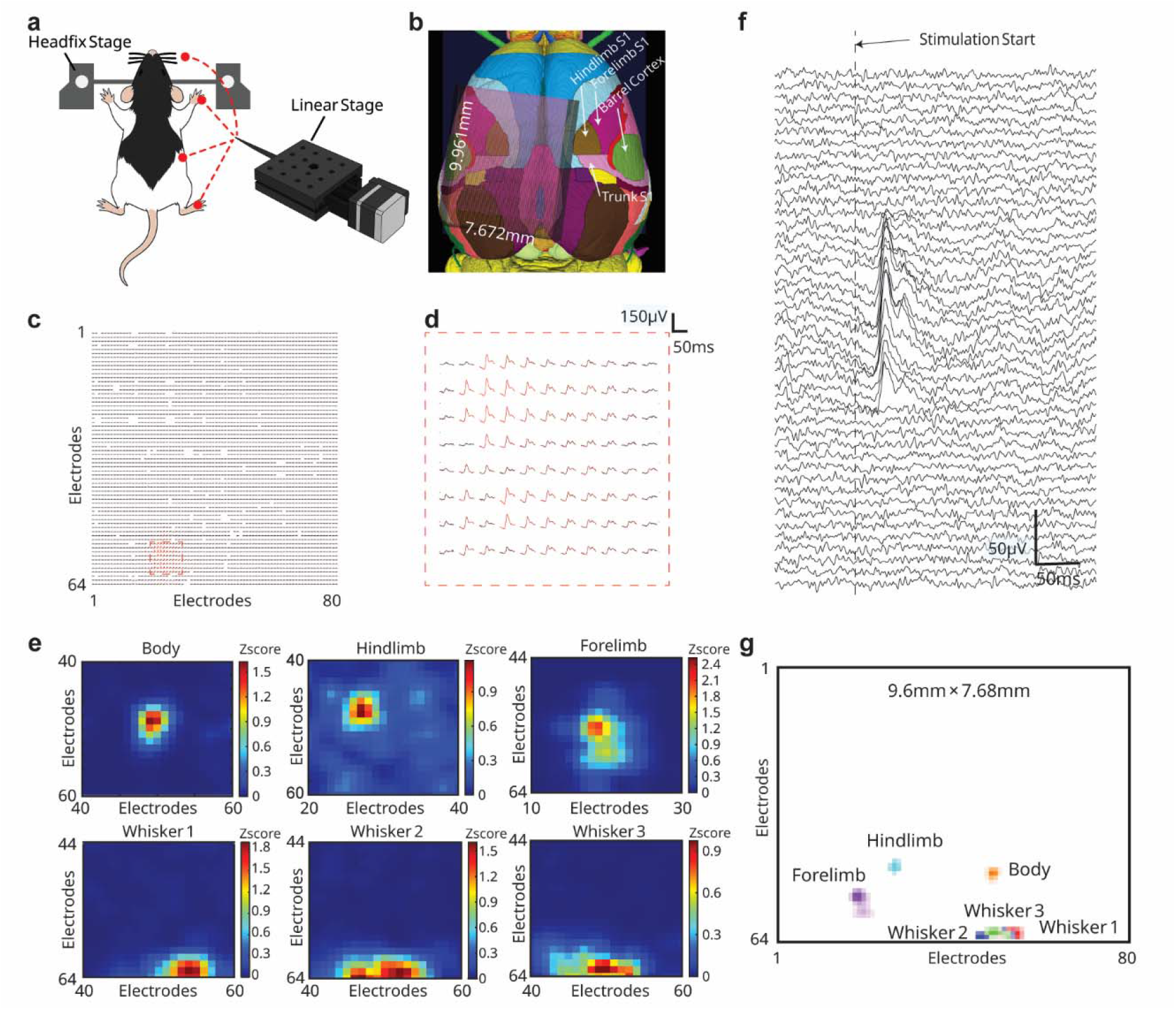
In vivo mapping of somatosensory cortical responses using the 5,376-channel μECoG platform. **a** Experimental setup illustration. Sensory stimuli (e.g., 1 □ms pulse, 1 □Hz) were delivered to the rat’s C-row whiskers, forelimb, hindlimb, and trunk (body), while neural activity was recorded from the left hemisphere using the head-mounted μECoG system and streamed to a PC. **b** Schematic showing the spatial placement of the μECoG array on the cortical surface, covering the left somatosensory areas. The full array spans ~7.7 □mm × 9.9 □mm, enabling cortical mapping at a high resolution. **c** Example evoked response amplitude map from right forelimb stimulation (n □= □50 trials, averaged 30 □ms post-stimulus). A distinct “hot spot” is observed in the left forelimb somatosensory area. **d** Zoomed-in view of the boxed region in c, showing clear sub-millimeter spatial structure with native 120 □µm pixel spacing. **e** Root mean square (RMS) amplitude maps averaged over 50 trials for stimuli to six body regions: trunk (body), hindlimb, forelimb, and three distinct whisker locations. Each stimulus evokes localized activation in spatially distinct cortical regions, demonstrating non-overlapping somatotopy. **f** Raw neural traces from two subsets of adjacent electrodes show localized and temporally structured evoked responses, with differences in onset latency across spatial locations. **g** Composite map overlaying activation patterns from multiple stimulation locations reveals distinct, spatially separated somatosensory domains on the cortical surface.

Each stimulus type was repeated over 20–50 trials to ensure response consistency. Trial-averaged traces revealed well-localized evoked responses, which became clearly distinguishable after common-mode subtraction (Fig. □5c). A zoomed-in view (Fig. □5d) highlights the system’s ability to resolve spatial features at the native 120 □µm pitch, capturing sub-millimeter neural activation patterns.

To quantify spatial response profiles, we computed the root mean square (RMS) amplitudes around stimulus onset from −100ms to 200ms. Every 5 ms frame was analyzed to identify the timepoint with maximum activation response. Heatmaps across trials revealed distinct, non-overlapping activation zones corresponding to stimulation of whiskers, forelimb, hindlimb, and trunk (Fig. □5e). Whisker stimulation evoked highly focal responses in the barrel cortex with peak activity occurring 20–25 □ms post-stimulation and decaying by ~50 □ms. Hindlimb and trunk stimuli appeared later (~85– 90 □ms onset), consistent with longer conduction pathways via the spinal cord. Overlaid maps from different stimulus locations revealed spatially distinct somatosensory territories (Fig. □5g).

Representative raw traces from a subset of nearby electrodes confirmed stimulus-locked responses and showed clear temporal differences in onset and amplitude between adjacent channels (Fig. □5f). These recordings show distinct neural activity, characterized by highly localized responses (~1 mm spread) with peak-to-peak amplitudes up to 70 µV.

## Discussion

Large-scale, high-resolution neural interfaces are essential tools for understanding the spatial organization and temporal dynamics of brain activity. However, the development of such systems remains challenging due to competing constraints on channel count, noise performance, physical form factor, power consumption, and integration complexity.

In this work, we present the highest-channel-count fully integrated, head-mounted μECoG platform to date, supporting simultaneous recording from 5,376 channels with on-chip stimulation, competitive electrical performance, and a compact 2.8 □cm × □2.5 □cm footprint (Table □S1).

Several key design considerations guided the development of this system. First, the platform was designed to capture the full bandwidth of neural activity (2 Hz–10 kHz), spanning LFP, AP, and ECoG. Second, probe and ASIC dimensions were pitch-matched to ensure scalability across different probe types. Third, multiplexing at the recording sites was deliberately avoided to minimize noise and crosstalk, prioritizing the fidelity of low-amplitude in vivo signals. Finally, we adopted a high-yield, post-processing-free bonding method, enabling cost-effective and scalable integration of ultra-dense flexible probes with custom ASICs.

In parallel with Neuralink’s efforts^38,39^, which focus on full-channel simultaneous AP recording and on-chip spike sorting, our work complements these efforts by enabling scalable, cost-efficient integration of flexible, ultra-dense probes with custom ASICs to support full-bandwidth raw neural data acquisition.

The system’s versatility was demonstrated in vivo. High-density cortical mapping in rats revealed precise spatial segregation of somatosensory fields, including whisker barrels, forelimb and hindlimb representations, and trunk responses. The ability to resolve these compact, functionally distinct regions, often just 1–2 □mm in size, highlights the importance of electrode density and uniformity in modern neuroscience tools. Importantly, our array achieves this without penetrating the brain, offering a minimally invasive approach suitable for chronic studies and potential human applications.

Beyond basic neuroscience, this platform opens doors for translational applications. In brain-machine interfaces, the combination of high spatial resolution and on-chip stimulation can support closed-loop neuromodulation and bidirectional interfaces. In clinical monitoring, such as epilepsy or motor recovery, large area μECoG arrays can provide richer spatiotemporal maps of cortical function than conventional grids. Moreover, the modular nature of the bonding interface allows integration with other probe types, including penetrating shanks, chemical sensors, and optoelectronic devices.

Looking ahead, future work will focus on incorporating wireless data transmission, real-time compression, and onboard machine learning to further reduce data volume and system footprint. Coupled with improvements in flexible electronics and biocompatible encapsulation, the platform presented here offers a scalable path toward next-generation neural interfaces for both research and clinical use.

## Methods

### ASIC design

The in-pixel LNA is implemented using an AC-coupled topology ^40,41,42,43^ (Figure. S1a) to eliminate input DC offset at the electrode. The LNA achieves a gain of ~34 dB using 12.5-pF input and 0.25-pF feedback capacitors. The input impedance is ~13 MΩ at 1 kHz, which is ~ 129 × higher than the probe impedance. A pseudo-resistor provides a low-frequency cutoff below 2 Hz to ensure coverage of LFPs ^43^. The LNA operates in weak inversion with a bias current of 2 µA to minimize the thermal noise. A fast recovery switch is integrated to reset the amplifier during and after stimulation, enabling artifact-free recording during high-rate stimulation protocols.

Following the LNA, a PGA provides tunable gain and filtering (Figure S1a,b). The PGA also uses AC coupling with tunable pseudo-resistors and incorporates a flip-over topology. This topology ^44, 45^ improves linearity compared to conventional PGA designs by symmetrizing the signal path and suppressing harmonic distortion. The low-frequency cutoff of the PGA stage is programmable via a control voltage applied to the pseudo-resistor, enabling flexibility across different recording modalities and experimental conditions. Frequency responses across different tunings are shown in Supplementary Figure. S1b. The embedded filter avoids the need for an additional explicit filter, minimizing the area within the pixel.

The chip integrates 32 independent stimulation units, each comprising a segmented current DAC and an H-bridge output stage for biphasic current delivery^46^. The H-bridge includes a dedicated active charge-balancing path that is enabled at the end of each biphasic pulse to neutralize any residual charge on the electrode-tissue interface. Stimulus timing and waveforms are controlled via the on-chip SRAM and an external FPGA. The stimulator’s reliability was validated by delivering 10 million biphasic pulses. The resulting current traces demonstrated near-perfect overlap and the mean current amplitude remained stable throughout (Figure S2b,c).

To support 1.3 Gb/s data throughput, the chip utilizes a dual-lane 12-bit Digital Video Port (DVP) interface (Figure S3), operating at 60 MHz clock per lane with offset synchronization. Each group of four rows is multiplexed into one ADC and time-multiplexed with row identifiers for accurate data reconstruction. An I^2^C interface allows programmable control over biasing, gain, stimulation parameters, and operating modes. The chip is mounted on a 2.8 cm × 2.5 cm headstage PCB with a stacked design including an Artix-7 FPGA, routing board, and power regulation circuitry. Data and power are transmitted through a single microHDMI cable, enabling lightweight, flexible connections suitable for freely moving animal experiments. Custom firmware and GUI were developed based on the Trodes platform to support real-time visualization and data acquisition from all channels.

The measurement setup for evaluating ASIC performance is shown in Fig. 2c. An on-chip switch matrix routes signals from an integrated ACin test pin to each pixel input, facilitating characterization with external equipment. During normal recording, this pin is left floating. For noise measurements, the input of each pixel was sequentially shorted to the reference electrode. For gain measurements, the input of each pixel was sequentially connected to a sinusoidal signal from an external signal generator. To evaluate the analog frequency response, a representative channel was routed to an analog output pin and characterized using a dynamic signal analyzer. For all tests, the digital output data from all 5,376 channels were streamed to the FPGA and processed on a host computer. Stimulator performance was evaluated under resistive (10 kΩ), capacitive (10 nF), and physiological (1x PBS solution) loads.

Stimulation reliability was validated by delivering 10 million biphasic pulses and measuring the waveform consistency across trials. The current traces demonstrated near-perfect overlap (Figure S2b), and the mean current amplitude remained stable throughout (Figure S2c). To evaluate charge balancing, stimulation was delivered into a capacitive load, and the voltage waveform showed symmetrical charging and discharging phases with a charge-balancing interval at the end (Figure S2d).

Impedance measurement and electroplating functionalities are implemented using in-pixel switches. Impedance is measured by applying an AC current via a coupling capacitor and sensing the resulting voltage through the recording path. Electroplating is performed using a DC current source accessed through the same in-pixel switch. These dual-use paths enable both initial calibration and long-term reliability monitoring.

### Probe design and fabrication

Guided by routing-density that could be achieved without sacrificing lithography yield, we = distributed the total channel count across four layers, with each layer containing 1,344 channels. Alignment marks were printed on the sacrificial layer, and each layer incorporated 1,344 interconnect lines with a minimal linewidth of 2 µm and a spacing of 2.5 µm. This multilayer approach allowed the use of standard photolithography, providing a reliable pathway toward mass production. The total thickness of the μECoG device was designed to be approximately 13 µm. Two μECoG designs were developed: one for rodents and one for humans. The rodent design features 25 µm-diameter electrode contacts with a 120 µm pitch, while the human design features 60 µm-diameter contacts with a 360 µm pitch. Both designs contain 5,376 individual channels, covering areas of 7.6 mm × 10 mm and 22.9 mm × 30.5 mm, respectively.

The microfabrication process was adapted from previous literature on depth probe fabrication ^47^. A nickel sacrificial layer was first deposited to facilitate release of the probe at the final step. Multiple layers of polyimide were then spin-coated, followed by photolithography, metal deposition, and lift-off to form the multilayered structures. The top and bottom polyimide substrates, each 5 µm thick (PI 2611 from HD MicroSystems), provided insulation and mechanical support. The four interconnect layers in the middle, responsible for linking the BGA pads to the electrodes, consisted of a 5 nm Ti / 150 nm Au / 5 nm Ti stack. A 1 µm-thick polyimide layer was placed between interconnect layers to insulate and protect the deposited gold traces. To expose the electrode sites and BGA pads for electrical connections, a 100 nm Ti hard mask was used to perform oxygen plasma dry etching in an ICP etcher, selectively clearing the polyimide in these regions. Perfusion holes were also etched in the electrode area to prevent the accumulation of saline beneath the thin-film device. The electrode material was deposited using iridium oxide (IrOx) sputtering, followed by a lift-off process. The typical electrode stack consisted of 10 nm Cr / 100 nm Pt / 10 nm Cr / 300 nm IrOx. Finally, the entire device was released from the substrate by dissolving the Ni sacrificial layer, after which the top layer was protected using photoresist. The device was then rinsed in acetone and IPA to remove the remaining photoresist, completing the fabrication process.

### Integration and packaging

The connection between the device and the ASIC is achieved through high-density semiautomatic ball bonding. A K&S semiautomatic wire bonder was used to perform the first and second layers of ball bonding. In the probe design, each bonding pad includes a hole, and gold bumps connect the exposed gold on the device to the exposed aluminum finish on the chip through two rounds of ball bonding.

The process begins by heating the substrate to 210°C. A free-standing wire is formed at the end of the pillar, creating a free air ball (FAB) using a 1 mil gold wire with a 2.4 mil FAB. The gold ball is then bonded by bringing the capillary into contact with the surface and applying ultrasonic force to securely attach it. The bonding parameters are as follows: ultrasonic generator power (USG) of 130–150 mW, bonding time of 40–70 ms, bonding force of 25–45 mN, and a USG ramp of 50%. Alignment tolerance must be maintained within 5–10 µm; otherwise, the first-layer ball may be deposited directly onto the polyimide substrate, which contains embedded signal metal lines beneath the polyimide layer. The capillary then rises to a specified height, where the wire tail breaks, leaving a residual length for the next FAB or bond. This process is repeated for each pixel until all bumps are completed.

Following the first-layer ball placement, the released device is manually positioned onto the first-layer gold bumps, which serve as alignment anchors. An ESD tweezer is used to gently push the device into position until it is flat. The second-layer ball placement is then performed, requiring precise realignment with the actual layout for each bond, as the thin-film device is manually positioned and typically has a tolerance of 5–10 µm. After completing these steps, all 5,376 backend pads are successfully bonded.

Once bonding is complete, the device is removed from the heat source, and underfill epoxy (Sikadur) and silicone (SYLGARD 186) are applied to secure and protect the connections, followed by curing. Ground and reference wires are soldered, and an additional layer of silicone is applied for additional protection. To ensure the electronics remain secure during acute or intraoperative procedures, a customized 3D-printed protective cage was designed and manufactured using a ProJet MJP 2500 Series printer.

### Surgery process

We performed a whole-brain craniotomy in a rat model during an acute implantation procedure. Animals were kept under anesthesia throughout the procedure, initially induced with 3–5% isoflurane and maintained at 1–2%, depending on body weight. A 10 mm × 12 mm rectangular craniotomy window was created for implantation. The typical boundary locations ranged from ML −5 to 5 mm and AP −10 to 2 mm, exposing the motor, somatosensory, and visual cortices. Once the craniotomy was completed, the dura mater was carefully removed to allow direct placement of the ECoG device onto the brain surface. For proper electrical referencing, the reference wire was placed in the frontal brain region, while the grounding wire was positioned in the neck tissues. A Faraday cage was then set up around the surgical area and grounded to the earth to minimize environmental noise and electrical interference. All surgical and experimental procedures were approved by the Institutional Animal Care and Use Committee (IACUC) at Rice University and comply with the National Institutes of Health guide for the care and use of Laboratory animals.

### Brain mapping experiments

We covered a substantial portion of the left hemisphere of the rat. After placing the ECoG, a resorbable hemostatic sponge (Goodwill) soaked in saline was placed over the craniotomy to ensure conformal contact between the probe and the brain surface. During the experiment, stimulation was applied at various locations using tweezers mounted on a motorized translation stage. Targeted areas included the body, hindlimb, forelimb, and specific whiskers. Each stimulation was repeated for 20 trials to ensure data reliability. The trial structure consisted of single-pulse mechanical stimulations delivered at a frequency of 1 Hz. The motor had a movable range of 5 mm and was programmed to complete each movement cycle within 100 ms. Neural recordings following each stimulation captured the evoked responses, and the results from all 20 trials were averaged to identify the stimulation effects.

### Data processing

To analyze the data, a 1–250 Hz filter was applied to isolate the ECoG frequency band, along with a 60 Hz notch filter, followed by root mean square (RMS) processing in 5 ms intervals. This enabled the creation of spatial heat maps of neural activity across all 5,376 pixels. First, the gain for each pixel was pre-tested to map the functional channels. Recordings from 2,688 channels from DVP1 and 2,688 channels from DVP2 were imported, totaling 5,376 channels, along with stimulation trigger signal timestamps extracted from the .h5 file. Common-mode noise was subtracted using signals from pixels located far from the tactile sensation and motor response regions. Baseline activities were identified from time frames with low RMS values, and these selected pixels were averaged and saved. After identifying trial start timestamps, we analyzed 200 ms before stimulation onset and 400 ms after onset, applied the channel map, subtracted common-mode noise, and downsampled the data to 1 kHz. The downsampled data was saved in a .mat file and stored as a 4-D array (dimensions: trial, column, row, time) for each trial. The RMS results were then plotted, and RMS averaging was applied to identify minimal differences between frames spaced 5 ms apart. Trials were averaged to quantify the statistical significance of signal localization, reduce background noise, and improve signal-to-noise ratio (SNR). Trials were aligned to the stimulus onset, and RMS values were plotted for each 5 ms interval. In the reduced pixel group, neighboring 4, 16, or 64 channels were combined, averaging their pixel amplitudes as the experimental group, while the original layout was used as the comparison group. The Pearson correlation coefficient was calculated between each pair of functional channels if both had gains within the specified range.

### Electrochemical analysis

The charge injection and storage capacities were evaluated using a Gamry Reference 600+ (Gamry Instruments, Warminster, PA). Measurements were conducted in a three-electrode setup with a large-area platinum counter electrode and an Ag/AgCl (3M NaCl) reference electrode (BASi Research Products, West Lafayette, IN). Voltage transients were recorded in response to biphasic pulses with a 100 ms pulse width and a 33 ms interphase interval at various amplitudes ranging from 3 µA to 18 µA. Cyclic voltammetry (CV) was performed with a sweep rate of 100 mV/s, scanning between 1 V and −0.8 V. Electrochemical impedance spectroscopy (EIS) measurements were conducted over a frequency range of 10 to 10,000 Hz.

### Bonding force experiment

We tested the bonding force using a Stellar 4000 Bondtester (Nordson) by applying a shear force to the bonded gold balls on the substrate. The shear force cartridge moved at 100 µm/s with a shear height of 5 µm. As the cartridge applied force to break the bonded balls, the relationships between time, forward displacement, and shear force were recorded. The shear force reached its maximum just before the gold balls were sheared off the substrate, and this maximum load was recorded. The experiment was repeated 10 times, and the maximum shear force was measured in each trial to determine the bonding strength. Both first-layer and second-layer bonding forces were assessed by controlling the cartridge head to shear off the first-layer and second-layer gold balls separately.

## Author contributions

Y.F. and Y.M. designed the hardware. Mattias Karlsson and Magnus Karlsson developed the FPGA firmware and data collection software. Y.F. and Y.M. performed the experiments and analyzed the data. C.X. and T.C. supervised the research. Y.F. and Y.M. wrote the initial manuscript and prepared the figures. P.Z., G.T., W.W., L.L., T.C., and C.X. revised the manuscript. All authors reviewed and approved the final manuscript.

## Data Availability

The datasets generated and/or analyzed during the study are available from the corresponding author on reasonable request.

## Ethics statement

All surgical and experimental procedures were approved by the Institutional Animal Care and Use Committee (IACUC) at Rice University (protocol no. IACUC-22-200) and were conducted in accordance with the National Institutes of Health Guide for the Care and Use of Laboratory Animals.

