## Supplementary Material for "High-channel-count neural recording and stimulation platform with 5,376 simultaneous recording channels"

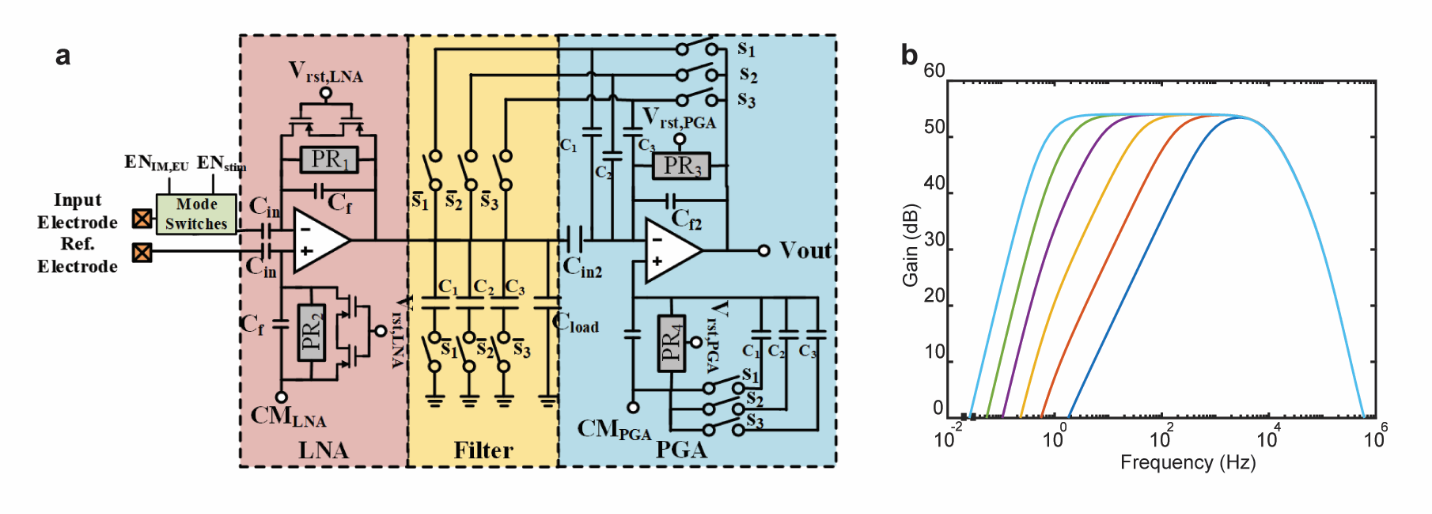

**Figure S1 Analog front end and simulation of tunable cut-off frequency**

a Circuit schematic of the analog front-end within each pixel, including the low-noise amplifier (LNA), embedded filter, and programmable gain amplifier (PGA).

b Measured frequency response of the recording channel under different low-frequency cutoff settings, achieved by tuning the control voltage of the pseudo-resistor in the PGA stage.

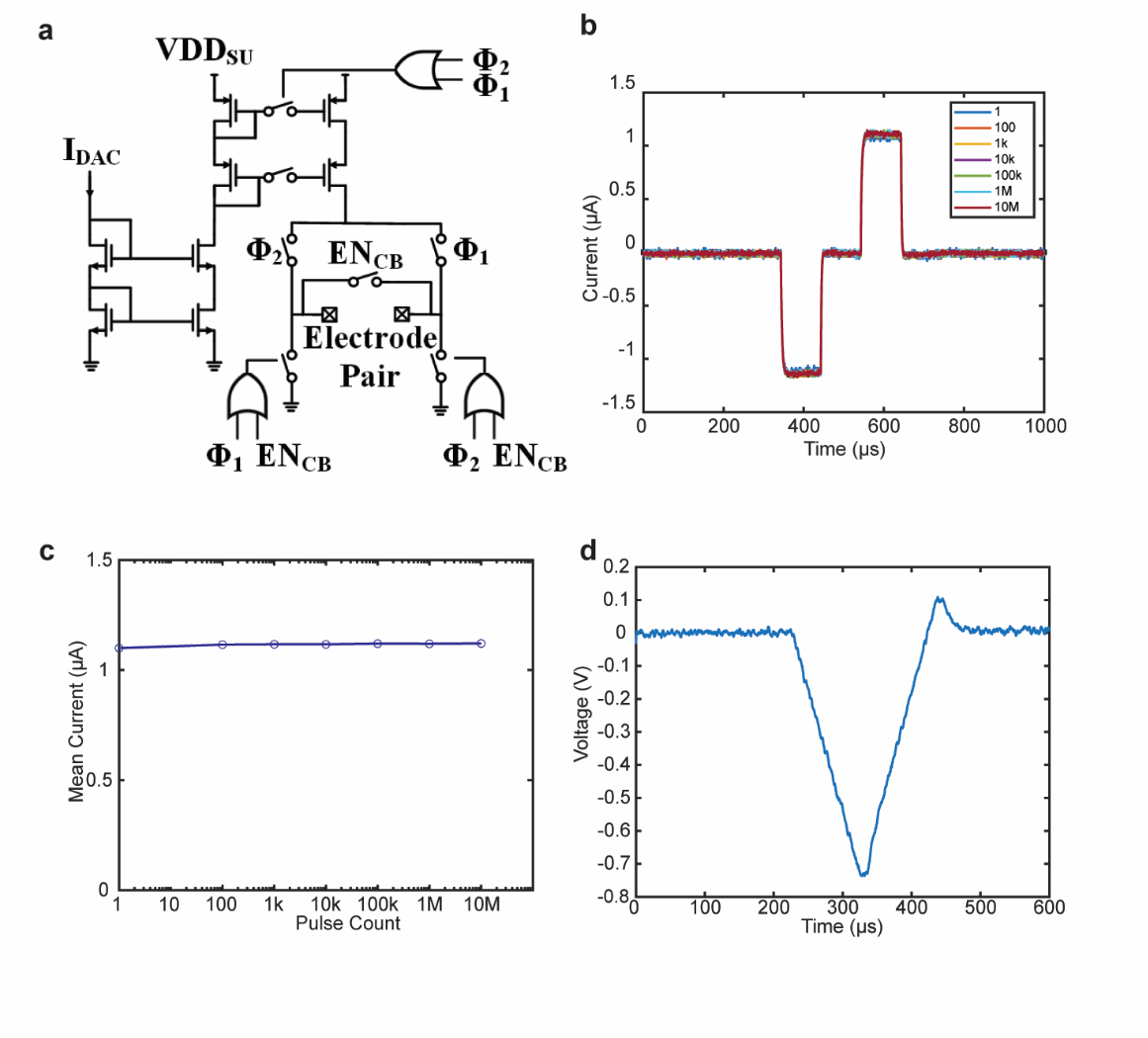

**Figure S2 Design and characterization of stimulation**

a Schematic of the H-bridge stimulator with integrated active charge-balancing switch (EN_CB_). The stimulation current is generated by a programmable current DAC (I_DAC_), which supports a range from 200 nA to 70 µA with 200 nA resolution using a well-matched current mirror structure.

b Long-term stimulation reliability test: 10 million biphasic pulses were delivered, showing tightly aligned waveforms across all trials.

c Mean current amplitude remains stable across pulse counts ranging from 1 to 10 million, validating long-term performance consistency.

d Stimulation response under capacitive loading, showing symmetric charging and discharging with a charge-balancing phase at the end of the pulse sequence.

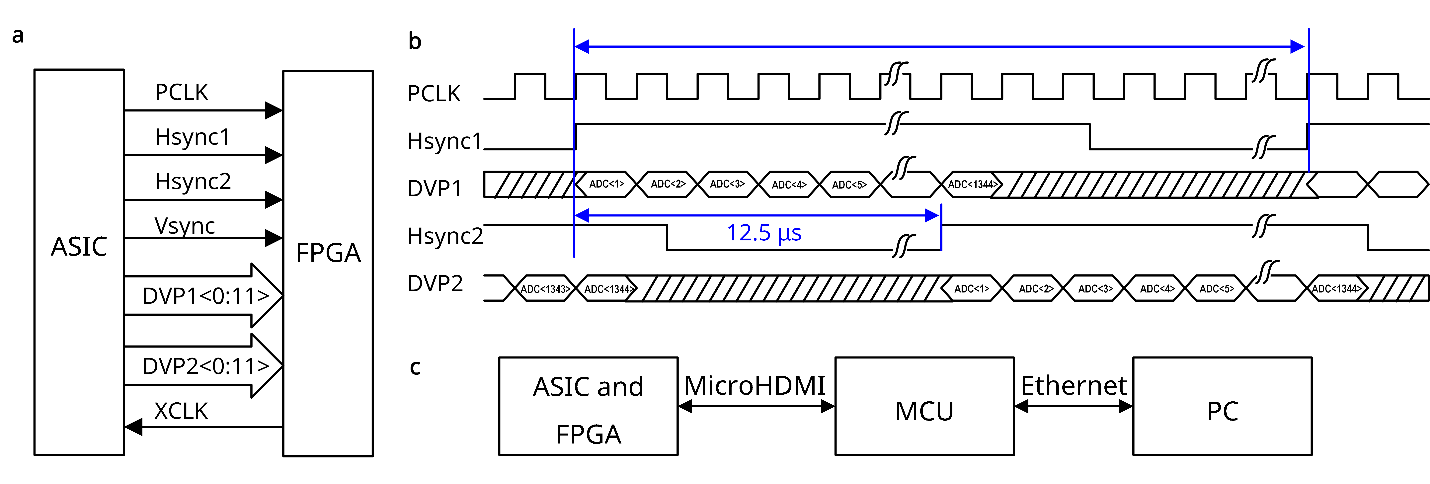

**Figure S3 Digital Video Port (DVP) protocol for high-speed neural data transfer.**

a Interface between ASIC and FPGA, showing pixel clock (PCLK), horizontal sync (Hsync1, Hsync2), vertical sync (Vsync), and two 12-bit parallel DVP data buses (DVP1<0:11>, DVP2<0:11>). The external clock (XCLK) synchronizes the system.

b Timing diagram of the DVP signals. Each Hsync triggers the transmission of a complete ADC data set over the corresponding DVP lane. In each 12.5 µs period, 1,344 ADC samples are serialized per lane, synchronized to PCLK.

c Data transmission chain from the headstage (ASIC + FPGA) through MicroHDMI to MCU, then via Ethernet to the PC for real-time data acquisition.

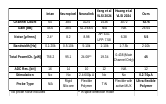

**Table S1 Comparison with other high-channel-count neural recording systems**^1,2^**.**

**Supplementary Section 1**

Our 5,376-channel neural recording system has a total power consumption of approximately 242 mW, averaging 45 µW per channel. A power breakdown (Figure S4) reveals that data transmission (I/O) is the dominant contributor at 34.2 µW/ch, accounting for over 75% of the total power. In contrast, the analog front end and digital backend consume only 6.7 µW/ch and 4.1 µW/ch, respectively. This high I/O power is due to the DVP interface (Figure S3) used for off-chip data transmission, which was selected for its straightforward integration with FPGAs and avoidance of clock recovery overhead. To reduce power in future implementations, we could incorporate data compression^3^ at the expense of raw data fidelity, or transition to a low-swing transmission protocol, which could reduce energy per bit by an order of magnitude^4^.

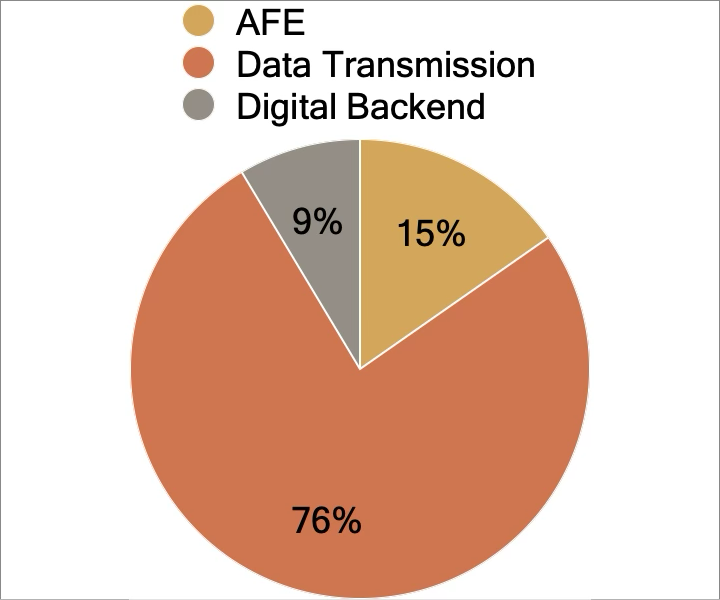

**Figure S4 Power breakdown of the system**
